# Sex and age specific effects on the development of addiction and compulsive-like drinking in rats

**DOI:** 10.1101/2022.12.28.522130

**Authors:** Jerome C. Foo, Ivan Skorodumov, Rainer Spanagel, Marcus W. Meinhardt

**Affiliations:** Department of Genetic Epidemiology in Psychiatry, Central Institute of Mental Health, Medical Faculty Mannheim, University of Heidelberg, Mannheim, Germany; Institute for Psychopharmacology, Central Institute of Mental Health, Medical Faculty Mannheim, University of Heidelberg, Mannheim, Germany; Department of Molecular Neuroimaging, Central Institute of Mental Health, Medical Faculty Mannheim, University of Heidelberg, Mannheim, Germany

## Abstract

Biological factors are known to influence disease trajectories and treatment effectiveness in alcohol addiction. One important biological factor influencing disease dynamics in alcohol use disorder is sex, as preclinical and clinical evidence suggests that gender influences disease trajectories in subjects with alcohol use disorder. Another critical factor is age at first intoxicating drink, which has been identified as a risk factor for later alcohol binging. Preclinical research allows prospective monitoring of rodents throughout the lifespan, providing very detailed information that cannot be acquired in humans. Lifetime monitoring in rodents can be conducted under highly controlled conditions, during which one can systematically introduce multiple biological and environmental factors that impact behaviors of interest.

Here, we used the alcohol deprivation effect (ADE) rat model of alcohol addiction in a computerized drinkometer system to study disease dynamics and transition points of addictive behavior in cohorts of adolescent vs. adult as well as male vs. female rats in direct comparison. Early age of onset of drinking (postnatal day 40) in male rats had surprisingly little impact on the development of drinking behavior and compulsivity (quinine taste adulteration) when compared to rats that started drinking during early adulthood (postnatal day 72). However, female rats showed a stronger resistance to quinine taste adulteration after a history of ADEs, indicative of more pronounced compulsive behavior in females as compared to male rats. Specifically, female rats exhibit during quinine taste adulteration a higher frequency of drinking events and a larger access size, especially from the 20% alcohol solution, resulting in fast intoxication despite taste aversion in a relapse-like situation. Our results suggest that drinking patterns, and specifically solution preference as well as access size, provide a valuable indication into determining compulsive drinking. These findings provide a better understanding of sex and age factors involved in the development of drinking behavior that are important in preclinical development of models of addiction, drug development and exploration options for new treatments.

## Introduction

Excessive alcohol use can have serious effects on health and is a leading cause of preventable death worldwide ((NCCDPHP), 2022a). Alcohol is the most common substance for which addiction criteria are met. Addiction is characterized by craving, loss of control of amount of or frequency of use, compulsion to use and continued use despite adverse consequences(WHO, 2022). While some people can consume large amounts without coming to harm, others experience ongoing addiction-related problems.

Clinical and preclinical research has documented sexually dimorphic effects of alcohol exposure through life, and both developmental processes and treatment effectiveness may differ accordingly (Finn, 2020). Epidemiological investigations suggest that men have the higher 12 month (17.6%) and lifetime (36%) prevalences for AUD than women (10.4, 22.7% respectively)(Grant et al., 2015), but this may be due to opportunity rather than specific vulnerability(Becker & Hu, 2008; Van Etten, Neumark, & Anthony, 1999). Although men are more likely to drink alcohol and consume excessive amounts, in women who drink excessively, biological differences (e.g. body size, structure, chemistry, etc.) are thought to lead women to absorb more alcohol and take longer to metabolize it ((NCCDPHP), 2022b), leading to more immediate as well as longer lasting effects. Women become addicted to alcohol more rapidly and at lower doses and have a faster progression to dependence (Zilberman, Tavares, & el-Guebaly, 2003); research has identified higher risk of liver diseases, accelerated alcohol-related cognitive decline and shrinkage of the brain in women compared to men (Erol & Karpyak, 2015). Indeed, in recent years it has been suggested that the ‘gender gap’ in drinking is decreasing ((Keyes, Grant, & Hasin, 2008) and has almost closed (Stelander et al., 2021). While preclinical research has in large part used adult male animals in models of addiction and its treatment, there have been an increasing number of studies examining the development of alcohol drinking behavior in females, finding that females acquire self-administration of alcohol more rapidly and consume more alcohol, but have reduced severity of withdrawal symptoms than males, possibly tied to differential sex hormones ((Becker & Hu, 2008; Becker & Koob, 2016; Surakka et al., 2021)

Alcohol is widely used in youth and underage drinking is a serious problem, and early onset of drinking increases alcohol use in adulthood (Bonomo, 2005; Pitkänen, Lyyra, & Pulkkinen, 2005). Youth drink less often than adults but binge when they do; 90% of alcohol drinks consumed by youth are during binge drinking (NIAAA, 2022). This can have serious health consequences as adolescent alcohol use can cause long-term changes in brain function and also brain structure (Crews et al., 2019; Spear, 2018), and it has been reported that alcohol misuse is the leading cause of death for youth (15-24 yrs old, (Mokdad et al., 2016)). Human and rodent studies have both found that adolescent ethanol exposure results in deficits during adulthood, including cognitive impairments and altered development of gray and white matter (Crews et al., 2019; Spear, 2018). A facilitatory effect of adolescent exposure to alcohol on adult intake has been suggested in several studies (Amodeo et al., 2018; Spear & Swartzwelder, 2014) but there are also conflicting reports showing no increased consumption (Jury et al., 2017) or sensitivity to aversion resistance (Nentwig, Starr, Chandler, & Glover, 2019). There are also reports suggesting that age of drinking onset is not a strong predictor of prospective alcohol intake and relapse-like drinking (Siegmund, Vengeliene, Singer, & Spanagel, 2005).

Preclinical models of voluntary drug intake offer the ability to study the longitudinal development of addiction processes (Spanagel, 2017). Rats develop quickly and are considered sexually mature around 6 weeks of age; one day for a rat is the approximate equivalent of one month of human life (Sengupta, 2013). The alcohol deprivation effect (ADE) paradigm is a longitudinal model of alcohol addiction development in which rats are given voluntary access to different concentrations of alcohol (Vengeliene, Noori, & Spanagel, 2013). The experimental procedure consists of repeated deprivation and reintroduction phases; reintroduction phases approximate relapse after abstinence and over time, rats show addiction-like drinking patterns, developing increased preference for stronger solutions of alcohol. In addition, once addiction-like behavior has stabilized, rats also develop compulsive-like drinking (aversion resistance, e.g., taste adulteration of alcohol solutions using bitter quinine), which is one important measure of addiction (Timme et al., 2020; Vengeliene, Bilbao, & Spanagel, 2014).

Using modern high-resolution sensing and recording systems, it is possible to follow rats as they develop addiction-like behavior (Vengeliene et al., 2013), building profiles to understand the development of drinking behavior, response to treatment (Foo et al., 2019), and compulsive drinking at a greater level of detail (Foo, Meinhardt, Skorodumov, & Spanagel, 2022). There is a continued need for animal models to characterize compulsive alcohol intake towards the identification of mechanisms that promote pathological drinking (Hopf & Lesscher, 2014), and it is particularly important to extend this research to female and adolescent groups.

In the present study, we examined the longitudinally assessed drinking behavior of female and male (adult and adolescent) rats acquired via digital “drinkometer” as they underwent a four bottle (H2O, 5%, 10%, 20% Ethanol) free-choice ADE paradigm (5 cycles over ∼11 months) and a subsequent quinine challenge. The drinkometer system continuously measured alcohol consumption at high resolution; locomotor activity was also concurrently acquired. We characterized and compared how drinking and movement behavior in the different groups evolved over the experiment, taking a closer look at consumption patterns (e.g., breaking down consumption by solution strength) during a regular ADE and during the quinine challenge.

## Materials and Methods

### Animals

Drinking and locomotor data from n=30 adult male Wistar rats (PND =72) are reported in (Foo et al., 2022). Data from adult females (n = 14; PND 66) and adolescent males (n =16, PND 44) from the breeding colony at the CIMH were collected and included. The CIMH Wistar rat line was developed at the Max-Planck-Institute for Psychiatry in Munich (“Crl:WI(Han)” (RS:0001833)) and has been selectively bred at the CIMH Mannheim for a robust alcohol drinking and ADE phenotype for over 15 years. Adult females and adolescent males were housed in the same experimental room.

Rats were housed individually in standard rat cages (Eurostandard Type III; Ehret) dimensions: top (outside) 425 mm x 276 mm, bottom (inside) 390 mm x 230 mm, height: 153 mm, floor area: ∼820 cm^2^ with feeders at the cover and followed by a 5 cm raised area, on a 12h:12h light-dark cycle (lights on at 07:30). During the experiment, all rats had *ad libitum* access to standard laboratory rat food (LASQCdiet® Rod16-Auto, Lasvendi and soy-free) and tap water. Bottles hold 250ml of liquid with ball nipples. The light intensity is max 130 lux in the drinking room and 50 lux within every cage. The relative humidity in all rooms is 45-65% and room temperature is constant between 22-24°C. Bedding material in the cages will have steam-sterilized aspen wood (2-3mm, Abedd) with no environmental enrichment. Cages are changed once per week. Our health monitoring program checks on all FELASA-recommended pathogens together with an external CRO (mfd Diagnostics) at a three month interval. We use at least one bedding/food-and water sentinel rat/per holding room. In addition, we have an extended pathogen screen once per year. Experimental procedures were conducted according to the ethical guidelines for the care and use of laboratory animals. Experiments were approved by the local animal care committee (Regierungspräsidium Karlsruhe).

### ADE paradigm, drinkometer system, quinine challenge

The ADE paradigm was carried out as previously described, with drinking behavior recorded using a digital drinkometer system (Vengeliene et al., 2013). Rats first underwent a long-term (4 weeks) period of baseline voluntary alcohol consumption. During this time, they are presented with four bottles, containing three different concentrations of EtOH (5%, 10%, 20%, prepared using 96% EtOH; VWR international #83804.360 diluted to the correct concentration) and water. Positions of bottles were changed regularly to avoid place preference.

After the first baseline period, rats are deprived of alcohol for 2 weeks, which is followed by reintroduction of alcohol. Upon reintroduction, the alcohol deprivation effect (ADE), a robust increase in alcohol drinking as well as a shift to stronger solutions, is observed. This process, with successive 2 week deprivations and 4 week reintroduction periods, was repeated 4 more times, for a total of 5 ADE cycles.

In the earlier study in adult male rats (Foo et al., 2022), during ADE6 cycle, a preliminary quinine test of different concentrations (0.01, 0.03 and 0.05g/L) was conducted, finding that 0.05g/L was appropriate for use in the drinkometer system bottles and had a larger effect than 0.01 and 0.03g/L on reducing consumption during the ADE. A full quinine challenge was conducted during ADE7 using 0.05g/L for the first three days of reintroduction. In the adult females and adolescent males, given the previous results, no preliminary test was performed; the quinine challenge was then conducted using 0.05g/L during the first 3 days of reintroduction during ADE6.

### Data Collection

A computerized Drinkometer system (TSE Systems) which records liquid consumption by amount continuously was used to monitor drinking behavior. The system is implemented in standard rat home cages (Eurostandard Type III) and has four drinking stations to enable choice of solution. Each of these consists of a glass vessel containing a liquid, and a high precision sensor that detects the amount of liquid removed from the vessel. Special bottle caps are used to prevent evaporation and spills. Bottle weights are measured in 200ms steps and can be registered every second, and the system can detect volume changes as small as 0.01g. For these experiments, the sampling interval was set at 1 minute.

Locomotor activity was monitored using an activity detection sensor (Mouse-E-Motion, Infra-e-motion) mounted above each cage. These devices use an infrared sensor to detect rat body movements of min 1.5 cm at any position inside the cage at the resolution of 1 second. For these experiments, locomotor activity was registered in 5 minute bins.

### Data analysis

Data were examined over the whole experiment and at the individual ADE levels. Analyses were conducted in R v3.63, and IBM SPSS 27.

The weekly level was examined to investigate overall developmental patterns of drinking and movement behavior. Average daily drinking during each BL and ADE week was calculated. In the Adult Females and Adolescent Males, technical issues led to lost data during the 4th BL period. In the young males, drinking data from 4 rats was lost from ADE4 until the end of the experiment.

To examine group differences during the whole experiment (from BL1 to BL6), a random-intercepts mixed model was used. In this model, consumption was specified as the dependent variable. Group (female, male, adolescent male) and period were specified as fixed factors. Restricted maximum likelihood estimation was used, with a diagonal covariance structure, which assumes heterogeneous variance and no correlation between time points. Pairwise comparisons were examined using estimated marginal means with Bonferroni correction.

ADE1, ADE5 and ADE with quinine (ADEQ) were further considered as the periods of interest. The first ADE has previously been shown to be an initial transition towards addiction-like behaviour (Foo et al., 2017), while ADEs are thought to stabilize by the 5th ADE. ADEQ was examined to study compulsive-like drinking behavior. During these periods, magnitude of ADEs were quantified by comparing total consumption (i.e., sum of all solutions) on the first day of reintroduction (RE) to the average consumption on the last 3 days of baseline (BASE) using paired T-tests. Welch’s tests were used to compare groups to account for uneven sample sizes (variances). Pairwise multiple comparisons tests with Bonferroni correction were performed to compare groups (females, males, adolescent males).

Drinking profiles were calculated for each rat. Consumption (g/kg) of each solution (5%, 10% and 20%) was calculated and Total Consumption (g/kg) was calculated by summing the solutions. Furthermore, we calculated the average access size and frequency of consumption of the different solutions (during the first day of ADE).

During ADEQ, compulsive-like drinking was quantified using percent change versus baseline of the same ADE cycle. That is:

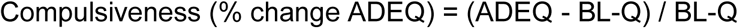

In addition, for an additional measure of compulsivity, comparing the size of the ADEQ to a regular one, we calculated the difference of magnitude in change:

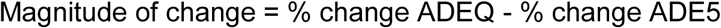

### Locomotor Activity

Locomotor activity was examined to characterize the different phases in a similar way to drinking data. Mean activity levels were calculated hourly and daily during the whole experiment and different periods. Association of hourly drinking by solution with locomotor activity was also examined. Mixed models were used to test for group differences as above.

In addition, we explored the circadian pattern of rats using a cosinor analysis. Cosinor analysis is often used to model circadian patterns in time series (Cornelissen, 2014). The package “ActCR” implemented in R was used to calculate the mesor (rhythm adjusted mean activity), amplitude (half the extent of predictable variation during a cycle) and acrophase (time to reach peak activity) of hourly locomotor activity during ADE1, ADE5, and ADEQ. For these analyses, 5 days of data (e.g., 120 hours) were used. (The first day of ADE until midnight was not used given the effect of the experimenter entering the room to change alcohol bottles.) For mesor, amplitude and acrophase, Welch’s tests were used to compare differences between groups with pairwise comparisons using Bonferroni correction.

## Results

### General drinking behavior between groups

The weekly total alcohol consumption is illustrated in **Figure 1a**, showing the average daily total consumption in different groups by week phase during the experiment. Significant effects of group (F(2,55.76501)=15.382, p < 0.001) and period (increased consumption over time, F(1,312.017)=53.194, p < 0.001) were observed. Adult female rats drank significantly higher amounts of alcohol than both adult male and adolescent male rats (both p < 0.001) throughout the duration of the experiment, descriptively starting even at the first BL. Adolescent males (starting drinking on PND44) and adult males (starting drinking on PND72) had similar drinking patterns throughout the entire experiment and consumption did not significantly differ (n.s.).

**Figure 1:**
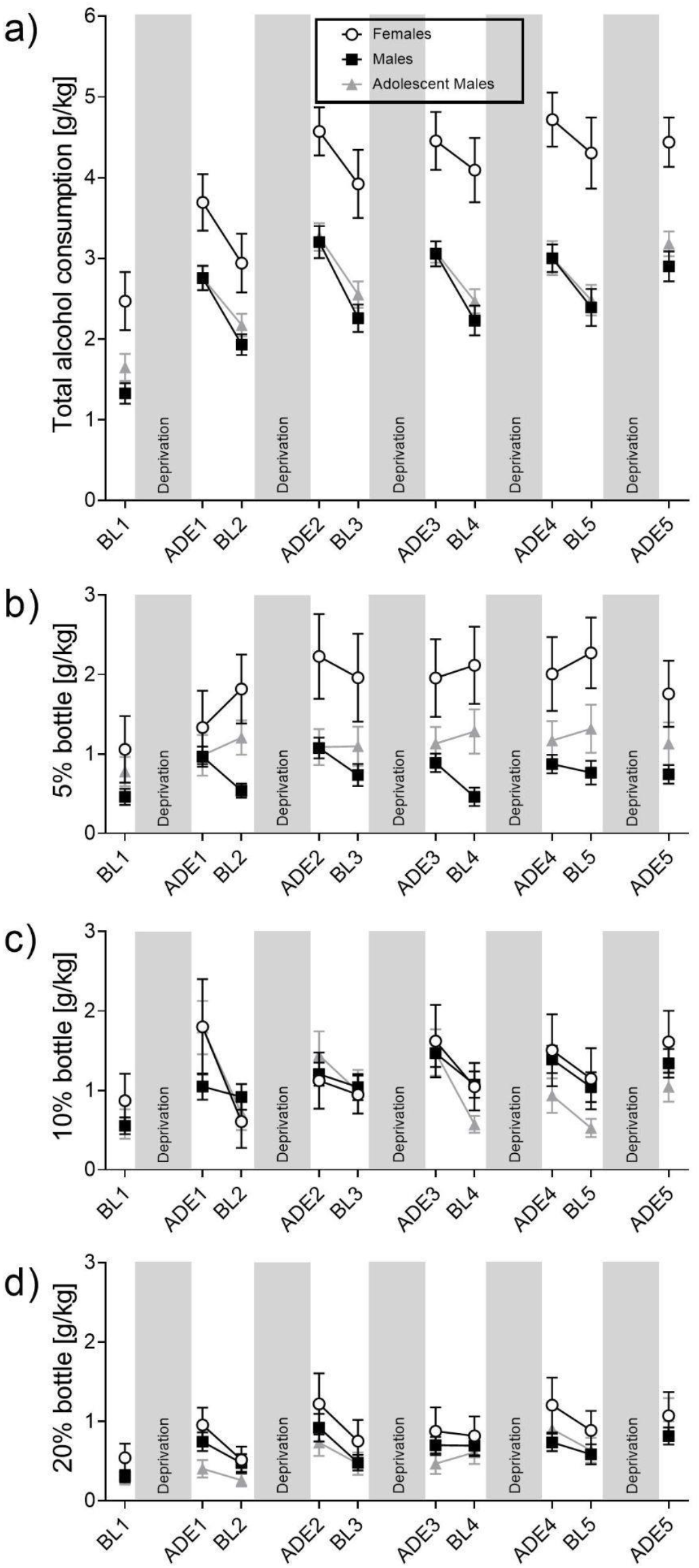
Drinking behavior between groups throughout the experiment. The total alcohol consumption (**a**) as well as individual bottles (5% -> **b**, 10% -> **c** and 20% -> **d**) are summarized for the different baseline (BL) and relapse (ADE) phases of the experiment. Data are presented as mean values ± SEM. Gray bars represent experimental deprivation phases, where only water was accessible.

Underlying this, as shown in **Figure 1b-d**, the higher levels of drinking observed in females compared to the other two groups was due to a significantly higher consumption of 5%. An effect of group on 5% drinking (F(2,56.798) = 8.044, p < 0.001) was observed. Females drank significantly more 5% than males (p < 0.001), and adolescent males (p = 0.038). Descriptively, an increase in 5% drinking in females was seen after the first ADE. 5% drinking between males did not significantly differ throughout the experiment (n.s.). An effect of period was not observed (n.s.). For both 10% and 20%, effects of period were observed (increased drinking over time, 10% (F(1,207.211) = 32.615, p < 0.001; 20% (F(1,229.385)=35.604, p < 0.001), but no effect of group (both n.s.)

### Comparison of ADEs over time

In ADE_1_, significant increases in intake (ADEs) were observed in all groups: females(t(13) = - 6.660, p < 0.001), males (t(29)=-10.116, p < 0.001) and adolescent males (t(15)=-12.095, p < 0.001) **(Fig 2a)**. Group differences were observed in both BL_1_ (Welch(2,27.685)=3.554, p = 0.042) and ADE_1_ (Welch(2,33.247)=4.117, p = 0.025), driven by differences between adult males and females (BL_1_: p = 0.007; ADE_1_, p = 0.038); drinking in adult and adolescent males did not significantly differ (n.s.).

**Figure 2.**
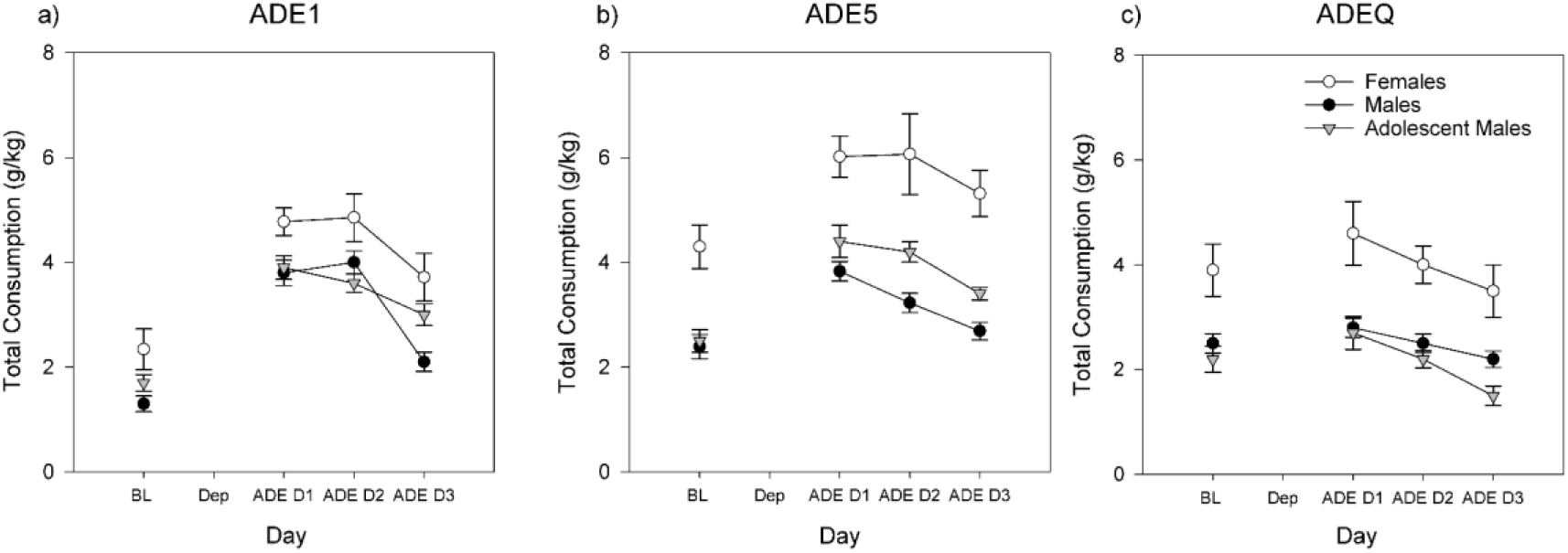
Total consumption of alcohol during: **a)** ADE_1_, **b)** ADE5 and **c)**ADEQ. Error bars denote SEM.

In ADE5, group differences continued to be observed (BL-5 Welch (2,20.952)= 8.647, p = 0.002; ADE5: Welch (2,16.565)= 12.181, p = 0.001) drinking in females was much higher than in adult (BL-5: p < 0.001; ADE5: p < 0.001) and adolescent males (BL-5: p=0.007, ADE5: p = 0.008)**(Fig 2b)**. BL and ADE drinking in adolescent and adult males did not significantly differ (n.s.). Significant ADEs were observed in all groups (all p < 0.001). We note that the average magnitude of ADEs was larger in males than females (M: +62%, AM: +76%, F: +38.5%), potentially due to a ceiling effect.

With the introduction of quinine as compulsivity measure during ADEQ **(Fig 2c)**, sizes of ADEs in all rats were decreased; only in adult males was a significant ADE (i.e., compulsive-like drinking) observed (t(29) = -2.289, p = 0.03; Adolescent males (t(10) = -1.899, p = 0.087; Females (t(13) = -1.033, p = 0.321), however the larger sample size plays a role in this statistical significance. As in other periods, group differences were observed (BL-Q: Welch(2,22.852)=4.655, p = 0.020; ADEQ: Welch(2,20.665)=3.995, p = 0.034) and the amount of drinking was higher in females than both males (BL-Q: p = 0.003; ADEQ: p = 0.001), and adolescent males (BL-Q: p = 0.004; ADEQ: p = 0.007), with no significant differences between adult and adolescent males observed (n.s.).

### Comparing regular and quinine ADEs by individual alcohol solutions

Differences between last regular relapse (ADE5) and the following quinine-adulterated relapse (ADEQ) are depicted in **Figure 3**. For the 5% bottle, while significant differences were observed during ADE5 (Welch(2,19.369)=3.858, p = 0.039), i.e., much higher consumption in females than males (p = 0.002) and adolescent males (p = 0.28) were observed. These differences between groups disappeared in ADEQ (Welch(2,18.814)=2.665, p = 0.173). Significant decreases between ADE5 and ADEQ were observed in all groups (adult males: t(28)=6.427, p < 0.001 ; adolescent males (t(9)=2.371, p = 0.042), females (t(14)=4.410,p < 0.001). For the 10% bottle, consumption decreased in all groups but this decrease did not reach significance (adult males: t(28)=2.028, p = 0.052, adolescent males: t(9) = 2.126, p = 0.062, females: t(14) = 0.325, p = 0.750). No significant group differences were observed.

**Figure 3.**
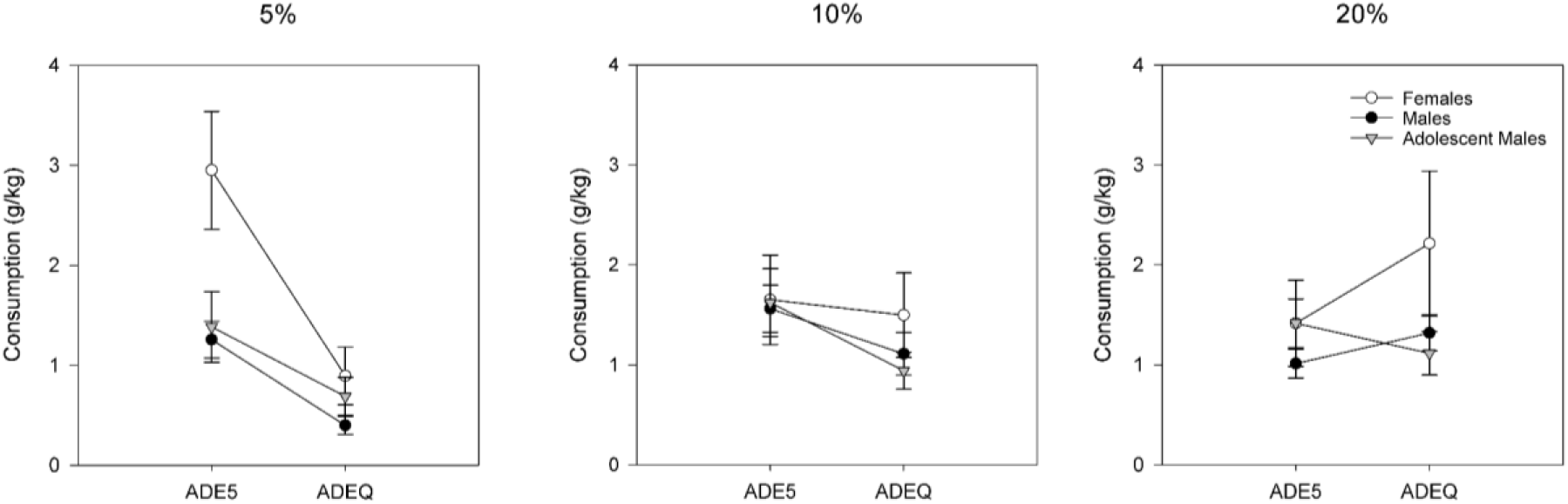
Differences in Consumption of 5%, 10% and 20% alcohol between the last regular ADE5 and ADE+ quinine (ADEQ). Error bars denote SEM.

For 20%, descriptively, consumption increases in females (the most pronounced, but with high variance) and adult males were observed; while a decrease was observed in adolescent males; none of these changes reached significance. No significant group differences were observed.

### Access sizes and frequency of accesses in regular ADE and ADEQ

Looking at ADE5 (**Fig 4a**), access sizes were largest in females for all solutions, especially for 20%. In ADEQ (**Fig 4b**), access sizes of 5% decreased in all animals compared to ADE5 (adult males: p < 0.001; adolescent males: p = 0.002; females: p = 0.028). Only adolescent males experienced a significant decrease in 10% access size (p = 0.002), and no group had significant decreases in 20% AS (adolescent males and females had non-significant increases).

**Figure 4.**
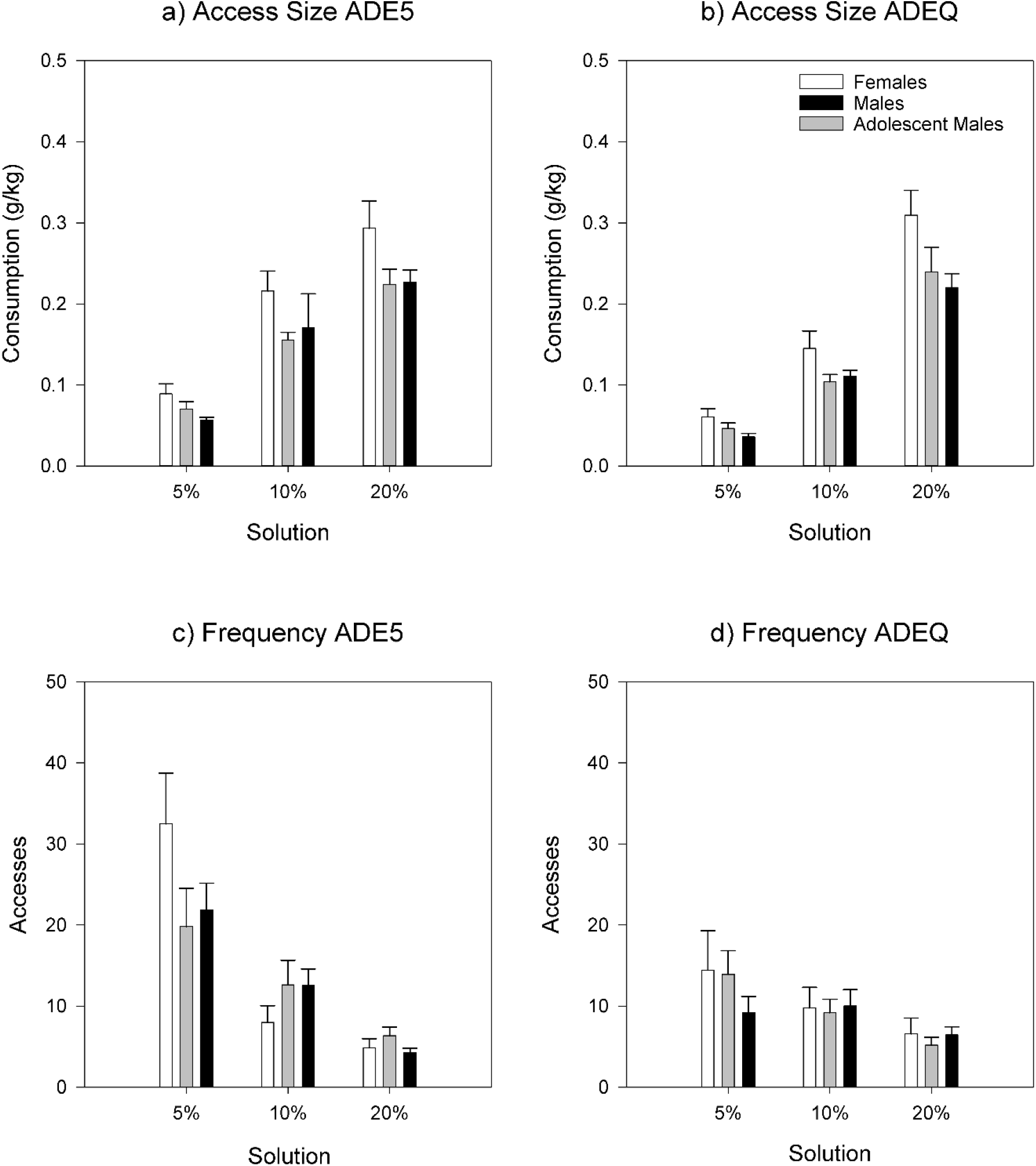
Access sizes for different alcohol solutions during **a)** ADE5 and **b)** ADEQ. and access frequencies for different alcohol solutions during **c)** ADE5 and **d)** ADEQ. Error Bars denote ±1 SEM.

In ADE5 **(Fig 4c)**, the frequency of 5% accesses was highest in females, while frequency of 10% accesses was higher in the male groups. Frequency of 20% accesses did not differ significantly between groups. In ADEQ **(Fig 4d)**, compared to ADE5, frequency of accesses was more affected than access size. Notably, the frequency of 5% accesses in females decreased almost 3 fold (p = 0.003), while 10% and 20% increased but not significantly. In adolescent males, frequencies of accesses to all solutions did not change significantly. In adult males, the frequency of 5% decreased (p = 0.003), while 10% decreased non-significantly; frequency of 20% accesses increased (p= 0.023).

### Compulsive-like drinking at the individual level

In order to assess the individual level of compulsive-like drinking in the three groups, we compared relapse rates of quinine-adulterated ADEs to baseline consumption at the same cycle, as well as the previous regular ADE. This painted a different picture of distributions of the different groups. Looking at %change during ADEQ, all groups had rats who exhibited compulsive-like drinking (i.e., positive change >0%; **Fig 5a**), and means did not significantly differ between groups (Welch (2,16.990) =0.969, p = 0.400), although we note high variability in the female group. Looking at the magnitude of change between ADEQ and ADE5, an effect of group was observed (**Fig 5b**, Welch (2,22.526)=5.747, p = 0.010), driven by females showing greater magnitude of change between ADEs than males (p = 0.003). Descriptively, using this metric, we also observe that male rats could be naturally broken into two “more” and “less” compulsive like groups.

**Figure 5:**
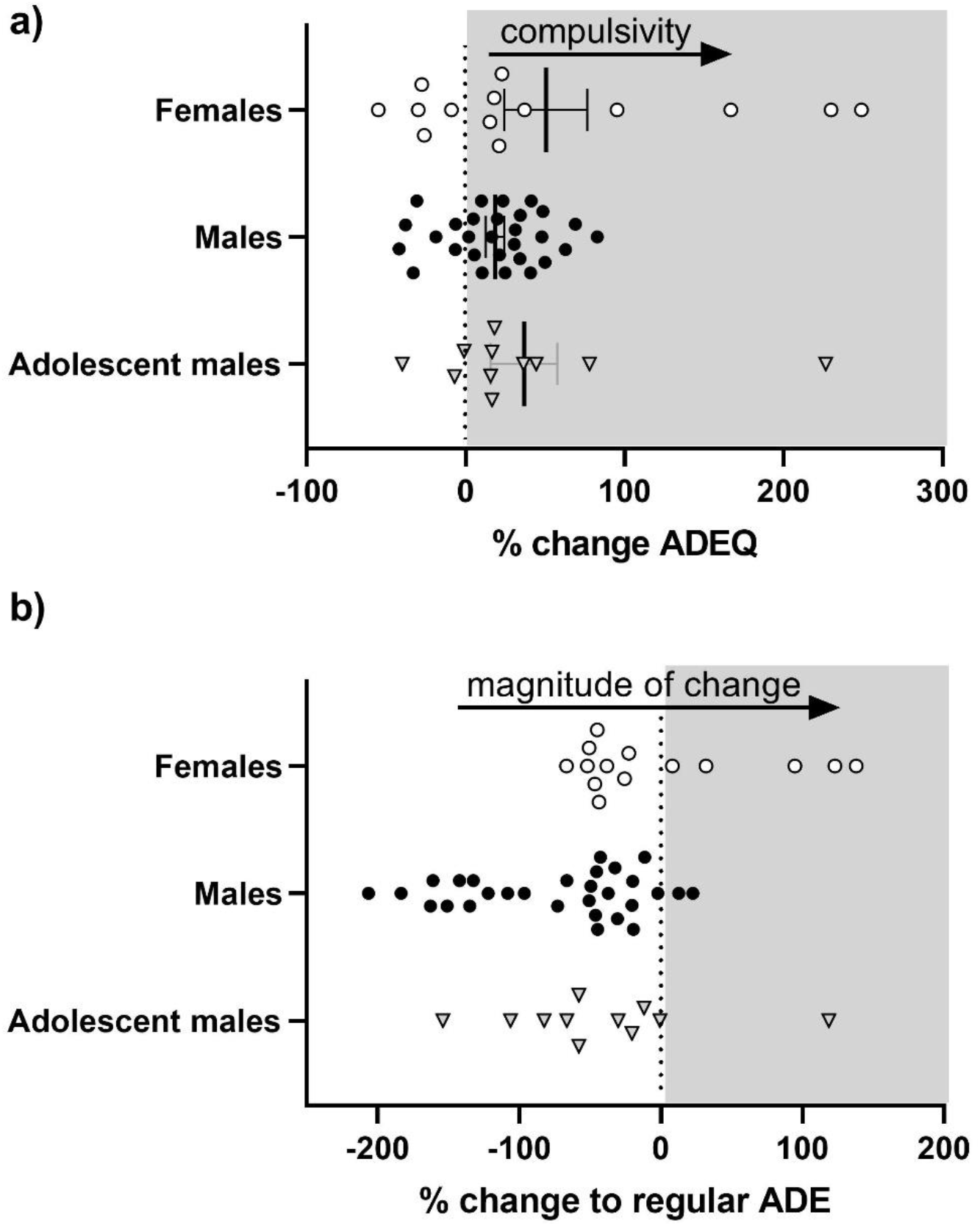
Compulsive-like drinking between different experimental groups. Compulsivity was calculated as **a)** % change ADEQ = (ADEQ - BL-Q) / BL-Q and **b)** the difference of magnitude in change: %change ADEQ = (ADEQ - BL-Q) / BL-Q - % change ADE5 = (ADE5 - BL-5) / BL-5.

### Overall locomotor activity

Average daily locomotor counts during the whole experimental period were higher in females, lower in adolescents, and lower still in adult males (Main effect of group F(2,54.769)= 31.759, p < 0.001; pairwise comparisons all p < 0.01). Over the whole experimental period, activity decreased in all groups (F(1,322.729)=81.530, p <0.001), descriptively, the most in adolescent males.

### Daily Movement Patterns

Examining hourly locomotor activity on the daily (**Figure 6**) and hourly (**Figure 7**) levels during ADE1, it was observed that females and adolescent males initially had similar movement patterns, while adult males moved less. By ADE5 movement in adolescent males had decreased to a level more similar to that in adult males; female activity levels remained comparatively constant.

**Figure 6.**
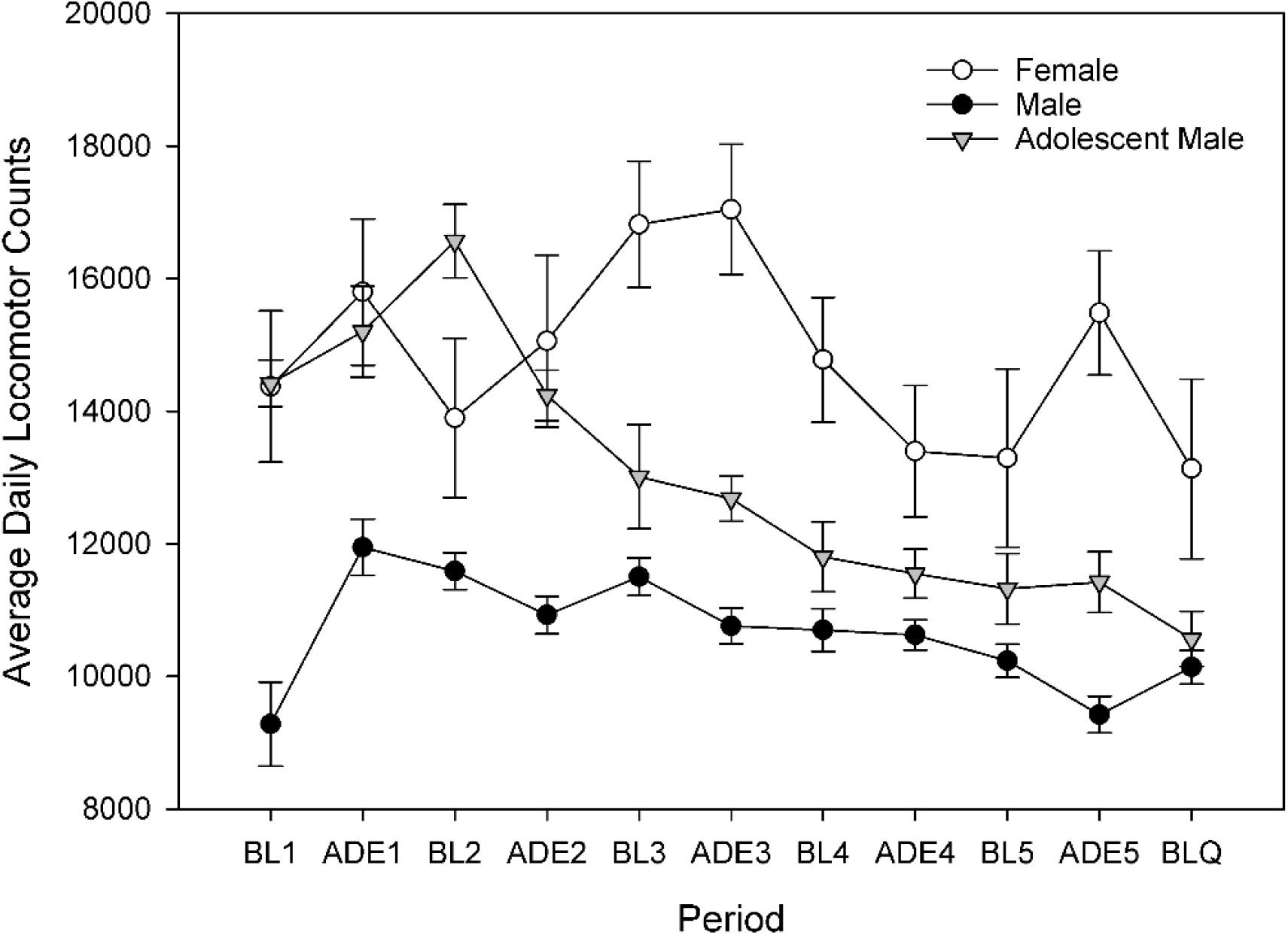
Average daily locomotor activity throughout experimental periods. Error bars denote ±1 SEM.

**Figure 7.**
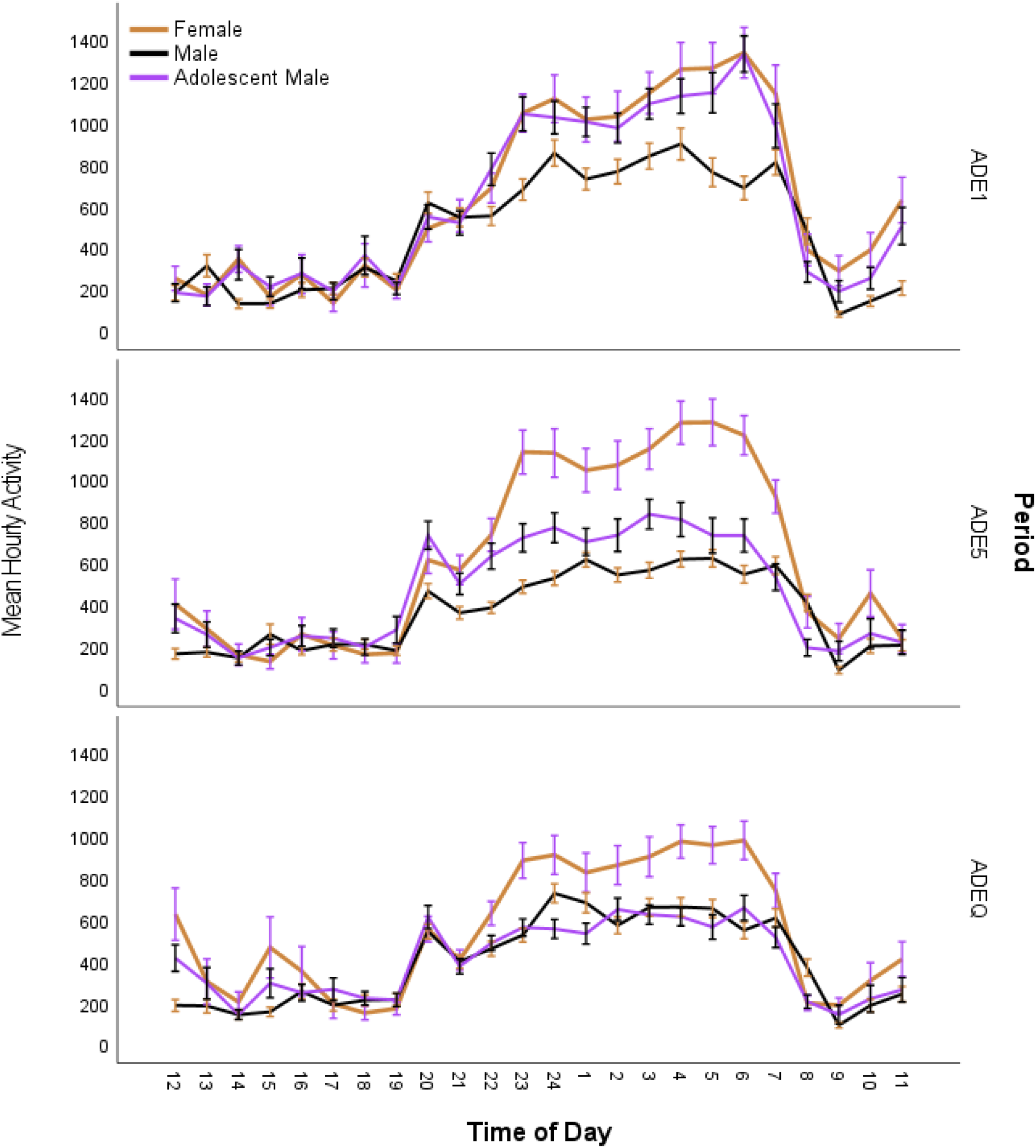
Mean hourly locomotor activity during the day in rats during different experimental phases. Error bars denote 95% CI.

During ADEQ, activity in both male groups was similar, and comparable to activity during ADE5. While still higher than males, female activity during ADEQ was reduced when compared to ADE5.

### Cosinor analysis

**Figure 8** shows the mesor, amplitude and acrophase as determined by cosinor analysis for the different periods. Significant group differences were observed in all measures during all periods (all p < 0.005). Table 1). Females had significantly higher mesor and amplitude than males during all periods (all p < 0.01), while adolescent males began the experiment with more movement rhythms to females and finished with more similar patterns to adult males. We note that the acrophase of female rats was later in the day than in both groups of males (ADE1: adult males p = 0.002; ADE5: adolescent males p = 0.001; ADEQ: both p < 0.05) this can also be seen in the hourly activity traces in **Figure 7**.

**Figure 8.**
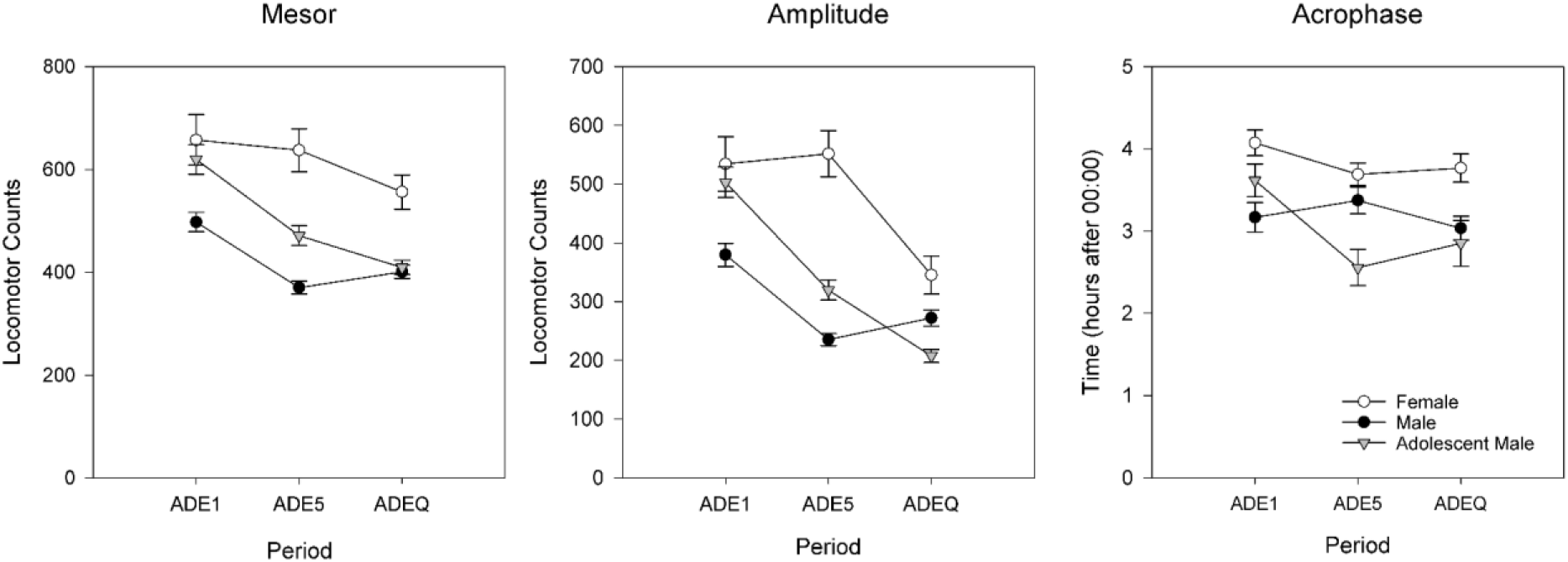
Differing circadian rhythm of rats over periods as modeled using cosinor: mesor (left), amplitude (center), acrophase (right). Error bars denote ±1 SEM.

## Discussion

In the present study, we investigated sex and age differences in the development of alcohol consumption and compulsive-like drinking. Our results point to considerably different male and female consumption patterns over time, differences which were also reflected in response to alcohol adulteration with quinine. Adolescent and adult males developed largely similar patterns of consumption.

Females consumed more alcohol than males during the whole experiment, which is in line with the wider literature. Interestingly, this difference was already apparent prior to the first ADE, which is consistent with the idea of the presence of different innate biological mechanisms promoting intake (Bell et al., 2003), and possibly the mediation of rewarding effects of alcohol by sex hormones (Torres, Walker, Beas, & O’Dell, 2014). While females consumed similar amounts of 10% and 20% as males, females consumed the 5% alcohol solution in much higher amounts. It is possible that this reflected taste preference; the 5% solution is slightly sweet and it has been shown that female rats prefer sweet solutions compared to male rats (Valenstein, Kakolewski, & Cox, 1967). Interestingly, the difference in 5% between males and females widened considerably after the first ADE cycle.

We observed that quinine decreased the consumption of 5% in all rats, with an especially large effect in female rats. Consumption of 10% was also decreased in all groups, whereas 20% consumption increased in adult animals but not adolescent males (an n.s. decrease). This suggests a possible age-related difference in the development of compulsive-like drinking; it could be that the compulsive-like drinking (and a shift to is not yet developed in younger rats, even as they show the similar consumption patterns to adult males during regular ADEs. In this study, we did not observe that adolescent male rats drank more alcohol or stronger alcohol than their adult male counterparts, which at first glance appears to come in contrast with literature finding that adolescent rats drink more per approach than adults and have more binge-like events (Bell et al., 2017; Dhaher, McConnell, Rodd, McBride, & Bell, 2012; Spear, 2015).

However, other studies report different findings (Nentwig et al., 2019) and these studies often have made observations using different paradigms (e.g., 2 bottle choice with 15% alcohol and water; providing different schedules of alcohol access, different routes of administration), or during more acute periods of time (e.g., 30 days) than in the present study, or with adolescent rats of different age; our results suggest that over (the life)time, the differences between (male) adult and adolescent rats may become less prominent.

One strength of the high-resolution multi-bottle approach is that it gives deeper insight into the mechanisms underlying behavioral effects of drugs and interventions. When comparing access sizes and frequency of accesses of specific solutions, we found differential patterns across groups and regular/quinine ADEs. Access sizes of all solutions were the largest in females. As previously shown (Foo et al., 2022), the effect of quinine was most apparent on the access size and frequency of accesses to the 5% solution. This effect was observed in all groups and especially prominent in the frequency of accesses in female rats; their original higher frequency of approaches 5% meant that they were the most affected. In contrast, in all animals 10% and 20% were less affected; only in adult males was there a significant increase in frequency of access to 20%. Looking at compulsive-like drinking at the individual level, we find that there is a wider variation of drinking patterns in females than the other groups. As a group, only adult males (with a larger sample) had statistically significant ADEs suggesting compulsive-like drinking; investigation of effects in a larger sample of female rats is warranted.

Our examination of movement patterns found similar results to drinking behavior, with larger differences observed between females and males than adolescent and adult males. Interestingly, the hourly data show that females and adolescent males started out with more similar movement patterns; while females maintained their activity levels over time, only decreasing mesor and amplitude with the introduction of quinine, both groups of males experienced reductions already during the regular ADE5. Another interesting observation was that female rats had later peaks of activity. Sex differences in circadian regulation (of sleep and waking cognition (Santhi et al., 2016) and circadian timing systems have been observed and are thought to stem from differences, for example, in (sex) hormonal and stress-responses (Bailey & Silver, 2014), neurophysiological (McCarthy, Arnold, Ball, Blaustein, & De Vries, 2012) and neuroendocrine (Mong & Cusmano, 2016) systems. Alterations in circadian timing are thought to be important in determining vulnerability to various types of disease, including addiction(Ehlers & Slawecki, 2000; Logan, Williams, & McClung, 2014; Rosenwasser, 2010) and circadian misalignment may disrupt reward mechanisms, promoting the transition from alcohol use to AUD (Hasler & Clark, 2013). One additional consideration when it comes to movement patterns is animal size. Females are smaller in size than males and cages are the same size regardless of animal size-this may have played a determining factor in movement patterns.

This experiment had certain limitations. First, females and adolescent males underwent their assessments at the same time, while adult males underwent the same procedure earlier. Although every effort was made to keep experimental factors constant, batch differences may have been introduced. Given the similarities observed between drinking patterns in the two male groups, however, this effect might be minimal. Next, it would be desirable to also include female adolescent rats in this design. Developmental differences in stress and gonadal sex hormones might be expected to introduce additional physiological and behavioral differences, (although it has been shown that estrous cycle phase does not account for all sex differences (Amodeo et al., 2018)). It has been previously suggested that female rats beginning drinking in adolescence may be more susceptible to stress-induced alcohol consumption (Füllgrabe, Vengeliene, & Spanagel, 2007). Also, sample sizes differed, with the adult males being the largest group; with equivalent sample sizes, statistically significant compulsive-like drinking might be more apparent in other groups.

We observed clear sex differences in drinking patterns, solution preference, response to aversive stimuli and movement patterns in male and female rats, suggesting potential areas of focus for future research. Gaining a more complete understanding of sex and age factors involved in the development of drinking behavior will be important in preclinical development of models of addiction, drug development and exploration options for treatment. Further refined exploration of these differences will be required to arrive at robust translational insights.

